# Interplay of target site architecture and miRNA abundance determine miRNA activity and specificity

**DOI:** 10.1101/214817

**Authors:** Giovanna Brancati, Sarah H. Carl, Helge Großhans

## Abstract

The recognition that the miRNA seed sequence is a major determinant of miRNA activity has greatly advanced the ability to predict miRNA targets. However, it has remained unclear to what extent miRNAs act redundantly when they are members of the same family and thus share a common seed. Using *in vivo* studies in *C. elegans*, we uncover features that drive specific target repression by individual miRNA family members. We find that seed-distal complementarity to a specific family member promotes specificity. However, the extent and robustness of specificity are greatly increased by seed match ‘imperfections’, such as bulges and G:U wobble base pairs. Depending on the seed match architecture, specificity may be overcome by increasing the levels of a miRNA lacking seed-distal complementarity. Hence, in contrast to a binary distinction between functional and non-functional target sites, our data support a model where functionality depends on a combination of target site quality and miRNA abundance. This emphasizes the importance of studying miRNAs under physiological conditions in their endogenous contexts.

## INTRODUCTION

MicroRNAs (miRNAs) are small RNAs of about 22 nucleotides that silence target messenger RNAs by binding to partially complementary sequences in their 3’ untranslated regions (3’UTRs). The miRNA loaded onto an Argonaute (Ago) protein forms the core of the miRNA-induced silencing complex that induces decay or translational repression of the targets (Krol et al., 2010). Conceptually, miRNAs can be separated into two main parts, the “seed”, comprising nucleotides two through eight, and the “seed-distal” 3’ end (Figure 1A). This is because the seed sequence has emerged as the main determinant for target identification (Bartel, 2009). Usually, functional miRNA targets contain “seed matches”, heptamers that base pair with perfect Watson-Crick complementarity to the miRNA seed. Systematic studies using ectopic miRNA expression found these seed matches to be necessary and sufficient for silencing (Brennecke et al., 2005; Doench and Sharp, 2004; Lai, 2002). Structural and biochemical analyses of the miRNA-induced-silencing complex (miRISC) have provided an explanation for the important function of seed matches: the seed of a miRNA bound by Ago exists in a pre-arranged conformation, thus reducing the entropic cost of binding and favoring duplex formation with a target (Chandradoss et al., 2015; Parker et al., 2009; Schirle et al., 2014).

**Figure 1.**
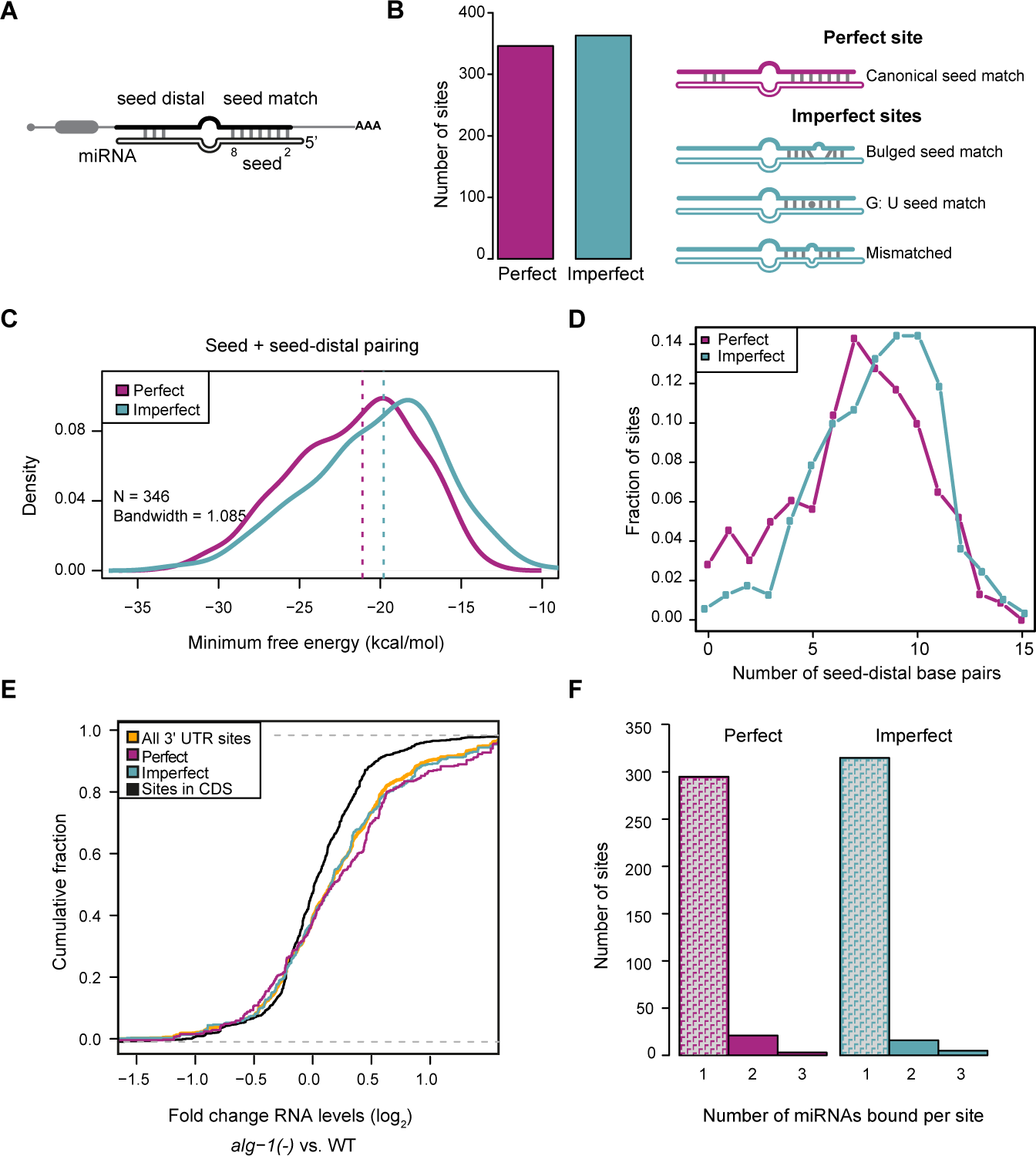
Sites with perfect and imperfect seed matches have similar features. A) Schematic drawing of a miRNA/target duplex with seed (nucleotides 2-8)/seed match and limited seed-distal pairing indicated. Top mRNA, bottom miRNA. B) Abundance of miRNA sites found in worm ALG-1 iCLIP chimeras (Broughton et al., 2016) that are located in 3’UTRs and bound by a miRNA belonging to a family. Sites are categorized by seed match quality. Perfect sites (magenta) contain 7 contiguous W-C base pairs from nucleotide 2 through 8. Imperfect sites (cyan) contain one mismatch in the sequence complementary to miRNA seed nucleotides 2-8, i.e., a bulge, a wobble base pair or another mismatch. C) Sites containing one seed mismatch have more extensive seed-distal pairing (number of nucleotides pairing beyond the seed) than perfect sites. D) Minimum free energy (MFE) calculation (done through RNAhybrid (Kruger and Rehmsmeier, 2006)) for miRNA:target site duplexes, as found in the chimeras. Imperfect sites median = −19.8 kcal/mol; perfect sites median = −21.1 kcal/mol). Dotted vertical lines indicate the median value for each curve. E) Upregulation of transcripts containing a perfect or imperfect miRNA binding site compared to sites in CDS (black, negative control) or any miRNA binding site located in 3’UTRs (yellow) in *alg-1(-)* relative to wild-type animals, based on RNA-seq data from Broughton et al. (2016). F) Number of miRNA sites bound by one, two or three members of a miRNA family and containing either a perfect or an imperfect seed match. We never observed more than three members bound to one site in this dataset.

miRNAs frequently occur in families, where family members share a seed sequence but differ in their seed-distal part. Given the reliance of target silencing on seed matches, it is assumed that miRNA family members can function redundantly, and, accordingly, widely used approaches predict miRNA targets for miRNA families rather than individual miRNAs (Bartel, 2009). Consequently, it was hypothesized that in order to attain specificity among family members, miRNAs require imperfect seed matches. In this scenario, an imperfect seed match would generally impair binding and activity of all family members, but extensive seed-distal base pairing would enable individual family members to compensate for the unfavorable binding (Brennecke et al., 2005).

Surprisingly, recently developed high-throughput biochemical methods that capture Ago-bound miRNA/target duplexes ligated into a chimeric molecule revealed interactions that frequently extended beyond the seed, to the seed-distal parts of the miRNA (Broughton et al., 2016; Grosswendt et al., 2014; Helwak et al., 2013; Moore et al., 2015). Moreover, in cell culture and *in vivo* assays, some targets were bound and silenced preferentially by specific family members, namely those that could base pair through their seed-distal parts (Broughton et al., 2016; Moore et al., 2015). Because such specificity also occurred for target sites with perfect seed matches, these findings argue that seed match imperfections are not a requirement for miRNA family member specificity.

Here we report that miRNA binding sites can predispose transcripts to silencing by a specific miRNA family member when it engages in extensive distal pairing. Although this occurs even in the presence of a perfect seed match, imperfect seed matches greatly enhance the extent and robustness of the effect. Moreover, specificity of targets with near-perfect seed matches can be overcome by miRNAs incapable of distal pairing when they are sufficiently abundant. Hence, although sequence-instructed, specificity is not fully hard-wired. These observations fit with a notion of miRNAs acting as rheostats on target mRNAs (Bartel and Chen, 2004), where the quality of the target site and the abundance of the miRNA act together to determine the regulatory outcome. Such a malleable mechanism for miRNA specificity not only expands the regulatory potential of miRNA families but also mandates that miRNA target validation occur under physiological conditions, in the absence of ectopic expression or overexpression of miRNAs. We illustrate both points by demonstrating that recoding *lin-41*, a specific target of the *let-7* miRNA, for regulation by all *let-7* family members, impairs normal *C. elegans* development.

## RESULTS

### Imperfect seed matches are abundant among functional miRNA targets but appear dispensable for miRNA specificity

Recent efforts to catalogue miRNA targets through Ago iCLIP (individual nucleotide-resolution cross-linking and immunoprecipitation) experiments have identified examples of targets that are regulated by only a single family member (Broughton et al., 2016; Moore et al., 2015). The finding that some of these sites contained perfect seed matches appear inconsistent with the canonical view that the presence of a perfect seed match equates with redundancy among family members (Bartel, 2009), but the role of seed match quality has not been investigated in the new studies. Hence, we sought to examine correlations between specificity and seed quality by re-analyzing published miRNA binding sites identified by sequencing of Ago iCLIP chimeric reads in *C. elegans* (Broughton et al., 2016). To this end, we filtered the reported miRNA/target chimeras located in 3’UTRs to specifically include only those that involved miRNAs that belonged to a family, which account for about 46% of the worm miRNAs (Agarwal et al., 2015). Subsequently, we divided the chimeras in two categories based on the seed match. “Perfect sites” contain a Watson-Crick match at position 2-8 of the miRNA seed, whereas “imperfect sites” contain a single bulge, a G: U wobble base pair, or a mismatch (Figure 1B).

Among the chimeras analyzed, we find imperfect sites to account for approximately half of the captured sites (346 perfect versus 363 imperfect chimeras in families, Figure 1B), and thus of much greater abundance than the < 5% expected from computational studies (Bartel, 2009). We observed that targets with imperfect seed matches tend to have an increased number of seed-distal matches to the miRNAs that bind them (Figure 1C), which may explain why duplex formation with these targets appears similarly energetically favorable as with targets that contain perfect seed matches (Figure 1D). Moreover, the analysis of RNA-seq data from wild-type and Argonaute mutant *alg-1(gk214)* (henceforth *alg-1(-)*) animals (Broughton et al., 2016), revealed that targets with imperfect sites in their 3’UTR are globally upregulated in *alg-1(-)* animals (Figure 1E), similarly to targets with perfect seed matches. With imperfect seed matches thus accounting for a substantial fraction of functional sites that have been captured by chimera sequencing, we sought to assess the impact of seed match quality on miRNA specificity. Surprisingly, we found that irrespective of the seed match, chimeras tend to be formed by only one specific family member (Figure 1F). In other words, an imperfect seed match appears dispensable for specific binding.

### *let-7* miRNA becomes dispensable for animal viability when the *lin-41* 3’UTR carries perfect *let-7* family seed match sites

Although the computational analysis suggested that extensive 3’ complementarity can suffice for miRNA-specific chimera formation, and presumably repression, it seemed possible that the observed miRNA/target specificity was driven partially or fully by non-redundant expression patterns among miRNA family members. Thus, a given target might be co-expressed with only one or a limited number of miRNA family members. Since the distribution of mature miRNAs across tissues is unknown for *C. elegans*, we cannot test for this possibility. Instead, we sought to examine experimentally whether and to what extent specificity relies on seed match quality and seed distal complementarity. We chose to focus our experiments on the *let-7* family because of its well-characterized mutant phenotypes, targets, and expression patterns.

The *let-7* family consists of *let-7* itself and its ‘sisters’ miR-48, miR-84, and miR-241 (Supplementary Figure S1A). *let-7* is required for animal viability because its absence causes de-repression of a specific target, *lin-41*, which in turn causes defects in vulval development and ultimately herniation of the gut through this organ (Ecsedi et al., 2015; Reinhart et al., 2000; Slack et al., 2000). This specificity in regulation by *let-7* contrasts with the fact that its sisters exhibit spatially and temporally overlapping expression profiles with *let-7* (Roush and Slack, 2008). Specificity was therefore hypothesized (Brennecke et al., 2005) to derive from the imperfect seed-matches in the two *let-7* miRNA Complementary Sites, LCS1 and LCS2, in the *lin-41* 3’UTR (Figure 2A and Supplementary Figure S1B, (Vella et al., 2004a; Vella et al., 2004b)). When bound by *let-7* family miRNAs, the seed match sequences of LCS1 and LCS2 generate an A bulge and a G:U wobble pair, respectively. Both sites also contain seed-distal complementarity to *let-7* but not its sisters.

**Figure 2.**
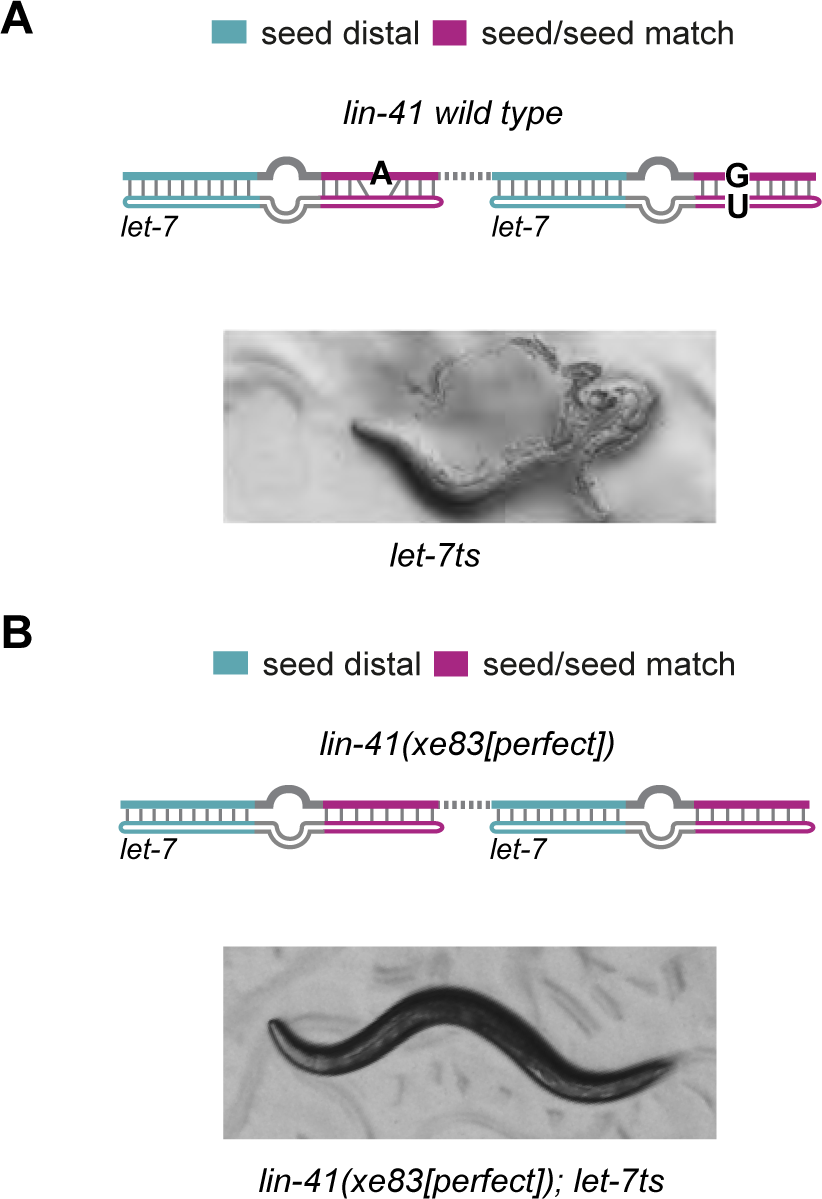
*let-7* becomes dispensable for viability when the *lin-41* 3’UTR contains perfect seed match sites. A), B) 100% of *let-7ts* mutant animals die at restrictive temperature (25°C) by bursting through the vulva as a consequence of the impaired *let-7/lin-41* interaction. (B) Lethality is suppressed when endogenous *lin-41* is altered to harbor binding sites with perfect seed match to the *let-7* family and unchanged seed-distal region. *lin-41(xe83[perfect]); let-7ts* double mutant animals survive and appear overtly wild type. *let-7ts: let-7(n2853) X*, temperature sensitive lesion. miRNA site legend: magenta = seed/seed match; cyan = *let-7* seed-distal binding.

To investigate whether seed mismatches are indeed required for specificity, we generated a new *lin-41* allele, *lin-41(xe83[perfect])*, with perfect seed matches to the *let-7* family in both sites (Figure 2B). We inactivated *let-7* by use of the *let-7(n2853)* temperature-sensitive mutant (henceforth *let-7ts*), which contains a change of nucleotide five in the *let-7* seed region and recapitulates the *let-7* null phenotype at the restrictive temperature, 25°C (Reinhart et al., 2000). Strikingly, whereas *let-7ts* single mutant animals died at 25°C by bursting through the vulva (Figure 2A), the two single point mutations in the 3’ UTR of *lin-41(xe83[perfect])* sufficed to overcome lethality with ≥98% (N=3, each with n≥200 animals) of *lin-41(xe83[perfect])*; *let-7ts* double mutant animals surviving into adulthood with an overtly wild-type appearance (Figure 2B).

### A perfect seed match allows redundant activity of the *let-7* sisters

To confirm that the perfect seed match of the *lin-41(xe83[perfect])* allele allows redundancy of the *let-7* family, we monitored the activity of the four miRNAs through a GFP reporter modified from (Ecsedi et al., 2015). In our assay, each animal contains a red mCherry reporter, which is used as reference during image analysis, and a GFP reporter, which is the miRNA activity sensor (Figure 3A). Both reporters are driven by the ubiquitous and constitutively active *dpy-30* promoter and contain the *unc-54* 3’UTR, generally thought to be devoid of regulatory elements. Finally, each reporter is integrated by Mos1-mediated single copy integration into a distinct genomic location (Frokjaer-Jensen et al., 2012).

**Figure 3.**
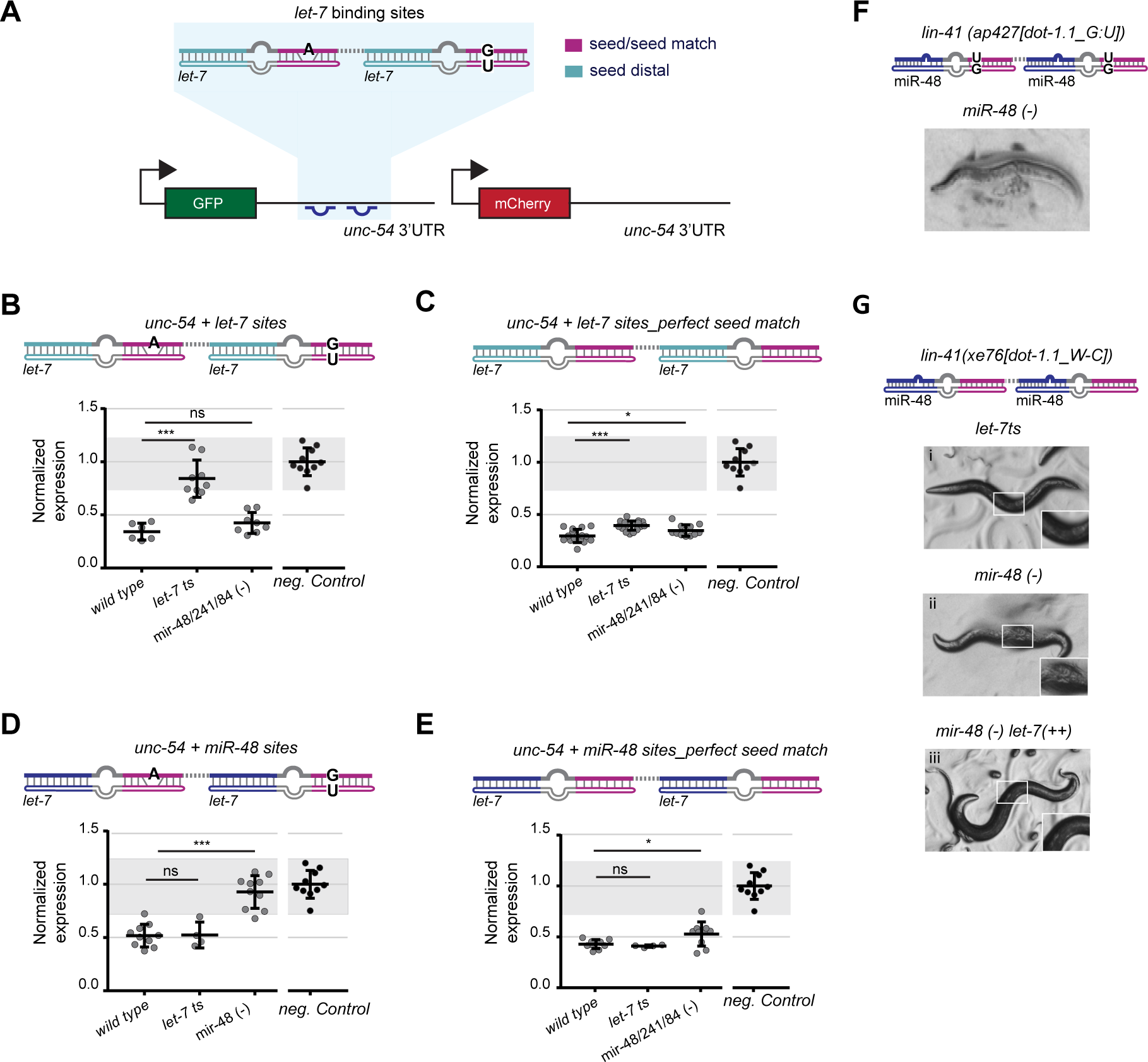
Imperfect seed matches and extensive 3’ pairing confer target specificity. A) Schematic of the reporters used to monitor miRNA activity *in vivo*. The GFP transgene *unc-54 + let-7 sites* reporter depicted contains 111 nucleotides of the *lin-41* 3’UTR (shaded in blue), which harbor the two *let-7* binding sites and their intervening sequence, grated into the heterologous *unc-54* 3’UTR. Transcription of the single-copy integrated reporter from the ubiquitously active *dpy-30* promoter is constitutive. Worms also contain a red mCherry reporter for normalization. miRNA site legend: magenta = seed/seed match; cyan = *let-7* seed-distal binding. B-E) Quantification of B) *unc-54 + let-7 sites* reporter, C) *unc-54 + let-7 sites_perfect seed match* reporter, D) *unc-54 + miR-48 sites* reporter, and E) *unc-54 + miR-48 sites_perfect seed match* reporter, respectively, in vulva cells of late L4 animals. Each dot represents the average of the GFP signal intensity, obtained by confocal imaging, divided by the mCherry intensity for a single animal per condition. 10-12 vulva cells were quantified per worm. Mean values are normalized to the average value of the GFP/mCherry ratio of the negative control *unc-54* 3’UTR reporter, which is not silenced. Horizontal line and error bars indicate mean values per condition ± SD. *P < 0.05 and ***P < 0.001, two-tailed unpaired t-test. For reference, data obtained for the *unc-54 (CTRL)* reporter are replotted in each of the four panels; gray shading is bounded by the min-max values of this control. F) Animals carrying the *lin-41(ap427[dot-1.1_G: U]* (Broughton et al., 2016)) allele die in the absence of miR-48. G) Converting the G:U wobble base pair at position 8 of both target sites generates strain *lin-41(xe76[dot-1.1_W-C]*, which survives loss of *let-7* (i) or miR-48 activity (ii), but becomes egg-laying (Egl) defective (93%, n = 132) in the latter condition. Overexpression of *let-7* (denoted as *let-7(++)*) animals suppresses the Egl phenotype of *lin-41(xe76[dot-1.1_W-C}]; mir-48(-)* animals (iii). *let-7ts: let-7(n2853) X, temperature sensi+ve* lesion, grown at the restrictive temperature 25 C*; mir-48(-): mir-48(n4097)V; mir-48/241/84 (-): mir-48/mir-241(ndf51) V, mir-84(n4037) X; unc-54(CTRL): wild type unc-54 3’UTR*.

To monitor *let-7* activity, we generated the reporter “*unc-54 + let-7 sites*” in which only a stretch of 111 nucleotides of the *lin-41* 3’UTR, comprising LCS1 and LCS2, were transplanted in the *unc-54* 3’UTR (Figure 3A). Silencing of this minimal target reporter by *let-7* was comparable to that of a reporter containing the full length *lin-41* 3’UTR (Figure 3B and Supplementary Figure S2A and B), confirming functionality. We focused our analysis on the vulva because *lin-41* repression by *let-7* in this organ is required and likely sufficient to prevent vulval rupturing (Ecsedi et al., 2015).

As expected, the “*unc-54* + *let-7* sites” reporter was expressed in young L1 or L2 animals (Supplementary Figure S2C), when the *let-7* family levels are low (Vadla et al., 2012), but extensively silenced in older, L4-stage larvae, when *let-7* family levels are high (Figure 3B). Moreover, it was de-silenced in *let-7ts* animals (Figure 3B), but not in animals lacking the three *let-7* sisters (*[mir-48/mir-241(ndf51)V, mir-84(n4037)X]*, henceforth *mir-48/241/84(-)*). Therefore, the stretch of 111 nucleotides suffices for efficient and specific *let-7*-dependent silencing.

Next, we generated an analogous *unc-54*-based reporter where the *let-7* complementary sites had perfect seed matches, “*unc-54 + let-7 sites_perfect seed match”*, as in the endogenous *lin-41(xe83[perfect])* mutation. Like the “*unc-54* + *let-7 sites*”, the new reporter was expressed in young L1 or L2 animals (Supplementary Figure S2C), but extensively silenced in L4-stage larvae (Figure 3C). However, unlike the “*unc-54* + *let-7* sites”, this new reporter remained strongly repressed in L4-stage larvae lacking *let-7* (*let-7ts*) or the three *let-7* sisters (*mir-48/241/84(-)*) (Figure 3C).

Taken together, the genetic interaction and the reporter assay data validate the hypothesis that the seed mismatches in the *let-7* complementarity sites of *lin-41* are necessary for specific regulation of *lin-41* by *let-7* proper to the apparent exclusion of the other family members.

### A seed-distal match establishes specificity to one miRNA in the presence of an imperfect seed match

Our observation that *lin-41* required an imperfect match to the *let-7* family seed to achieve regulation by only *let-7* appeared inconsistent with the results of the Ago iCLIP chimeric reads, which show preferential target binding by individual family members also in the presence of perfect seed matches. Thus, to challenge our finding, we sought to reprogram the LCSs to another *let-7* family member, miR-48, and test the effect of seed match imperfections. We chose miR-48 because its expression levels and spatial expression patterns appear very similar to that of *let-7* (Abbott et al., 2005; Martinez et al., 2008; Roush and Slack, 2008). We created an “*unc-54 + miR-48 sites*” reporter such that miR-48 would be capable of forming imperfect duplexes analogous to those formed by *let-7* on the “*unc-54* + *let-7 sites*” reporter (Supplementary Figure S3A). In agreement with our predictions, the “*unc-54 + miR-48 sites*” reporter was repressed at the L4 stage in both the presence and absence of *let-7* miRNA, but became de-repressed when miR-48 was absent (Figure 3D). Extensive seed distal complementarity was required for functionality: a reporter that base-paired only to nucleotides 13-16 of miR-48, chosen because structural data suggest that base-pairing between nucleotides 13-16 of the miRNA and a target may be favored (Schirle et al., 2014), failed to be silenced even in wild type conditions, i.e., with both *let-7* and miR-48 present (Supplementary Figure S3B).

Consistent with our results for the *let-7* reporters, the specificity of the “*unc-54 + miR-48 sites*” reporter was largely lost when we modified it to contain perfect seed matches: the resulting “*unc-54 + miR-48 sites_perfect seed match*” reporter continued to be silenced extensively in both *let7ts* and *let-7* sisters *mir-48/241/84(-)* animals (Figure 3E). However, silencing appeared marginally impaired in the absence of the *let-7* sisters (Figure 3E), mirroring an analogous result for the “*unc-54 + let-7sites_perfect seed match”* reporter in *let-7ts* animals (Figure 3C). We conclude that the imperfect seed match and the extensive 3’ pairing are both major determinants for the robust target specificity of the *lin-41* sites.

### A G:U wobble base-pair in a peripheral seed match location promotes miRNA specificity

The duplexes formed between *let-7* and *lin-41* contain a bulge between nucleotides 4-5 in LCS1 and a G: U wobble base-pair at position 6 in LCS2 (Figure 2A and Supplementary Figure S1B). We wondered if such centrally located ‘imperfections’ were required for specificity. We turned to the miRNA binding site in the *dot-1.1* 3’UTR, which had been shown to be specific to miR-48 (Broughton et al., 2016). Broughton and colleagues found that substitution of the *let-7* complementary sites in the endogenous *lin-41* 3’UTR by two copies of the *dot-1.1* site rendered animals insensitive to loss of *let-7* (Broughton et al., 2016), but made them depend on the presence of miR-48. This finding was attributed to the fact that the site features an extensive seed-distal match to miR-48 (Figure 3F and Supplementary Figure S3C). However, we noticed that the *let-7* family/*dot-1.1* predicted duplexes exhibited not only a perfect Watson-Crick pairing from nucleotides 2-7, but also a G:U wobble pair at position 8 (Supplementary Figure S3C). Although hexameric seed match sites complementary to nucleotides 2-7 are considered canonical and functional (Bartel, 2009), genome-wide studies also suggested that they are less functional than heptameric sites that match nucleotides 2-8 (Baek et al., 2008; Chandradoss et al., 2015; Grimson et al., 2007). Since G:U wobble base pairs elsewhere in seed-seed match duplexes appear detrimental to silencing (Brennecke et al., 2005; Didiano and Hobert, 2006; Doench and Sharp, 2004; Lai et al., 2005; Wolter et al., 2017), we wondered if this “peripheral G:U” in seed match position 8, might affect silencing and specificity.

To test this hypothesis, we modified the endogenous target sites in *lin-41* to those of *dot-1.1*, but with the G:U wobbles at positions 8 converted to Watson-Crick G:C pairs, yielding allele *lin-41(xe76[dot-1.1_W-C])* (Supplementary Figure S3D). We then compared the reliance of this and the *lin-41(ap427[dot-1.1_G:U])* strain, which carried the unmodified G:U-wobble-containing *dot-1.1* sites, on *let-7* and miR-48 for survival. Whereas both strains were insensitive to loss of *let-7* (Figure 3G(i) and (Broughton et al., 2016)), *lin-41(ap427[dot-1.1_G:U])* but not *lin-41(xe76[dot-1.1_W-C])* required miR-48 for survival into adulthood (Figure 3F and 3G(ii)). We conclude that the G:U wobble at position 8 repels binding by all *let-7* family members such that only miR-48, which can compensate through extensive complementarity of its 3’sequence, can exert repression. Collectively, our data thus reveal that bulges or wobbles in different positions of a seed match can serve to avoid redundancy of the *let-7* family and confer strong target specificity.

### miRNA abundance affects silencing *in vivo*

Although our experiments provided strong evidence that seed mismatches are required for robust specificity among *let-7* family members, we consistently observed evidence of residual specificity even for targets that contained a perfect seed match. This was true at the level of target reporters, where we observed modest but robust de-silencing specifically when the family member with seed-distal match was lost (Figure 3C, E), and phenotype (Figure 3G(ii)). Thus, although *lin-41(xe76[dot-1.1_W-C]); mir-48(-)* animals survived into adulthood, they exhibited an egg-laying (Egl) defect (Figure 3G(ii)) consistent with incomplete repression of *lin-41* (Ecsedi et al., 2015).

We wondered if this partial specificity could be overridden by increased levels of another miRNA family member. Since we were unable to overexpress mir-48, we tested this possibility by overexpressing *let-7*. *Mos1*-mediated single copy integration (Frokjaer-Jensen et al., 2014) of a genomic fragment, known to rescue *let-7* lethality (Reinhart et al., 2000), to a locus on chromosome V that is ~5cM apart from that of mir-48, yielded a ~2-fold increase in expression levels (data not shown). Consistent with our hypothesis, *lin-41(xe76[dot-1.1_W-C])* animals that over-expressed *let-7* were no longer Egl in the absence of miR-48 (Figure 3G(iii), compare to 3G(ii)). We conclude that *in vivo*, miRNA levels can affect silencing, and in particular override the specificity imparted by the seed-distal pairing.

### Seed match imperfections antagonize loss of specificity upon miRNA overexpression

Since the modest preferential silencing imposed by the seed-distal pairing to miR-48 could be overcome by increasing the levels of *let-7* in the presence of a perfect seed match (Figure 3G), we wondered about the effect of *let-7* over-expression on sites with more extensive target specificity. Therefore, we examined two reporters specific to miR-48 that harbored imperfect seed matches: the “*unc-54 + miR-48 sites*” (Figure 3D and 4A) and the “*unc-54 + dot-1.1 sites*” reporter, obtained by inserting two copies of the binding sites from the *dot-1.1* 3’UTR (Figure 4B). Consistent with the *in vivo* data ((Broughton et al., 2016) and Figure 3G), silencing of both reporters was dependent on miR-48 but not *let-7* (Figure 3D, 4A, 4B and Supplementary Figure S3E). By contrast, the response of the two reporters differed when we overexpressed *let-7* in the absence of miR-48: whereas the *unc-54 + miR-48 site* reporter (with central seed mismatches) was insensitive to a doubling of *let-7* expression (Figure 4A), silencing of the *unc-54 + dot-1.1 site* reporter (with peripheral seed mismatches) was restored to almost wild-type level in the same conditions (Figure 4B).

**Figure 4.**
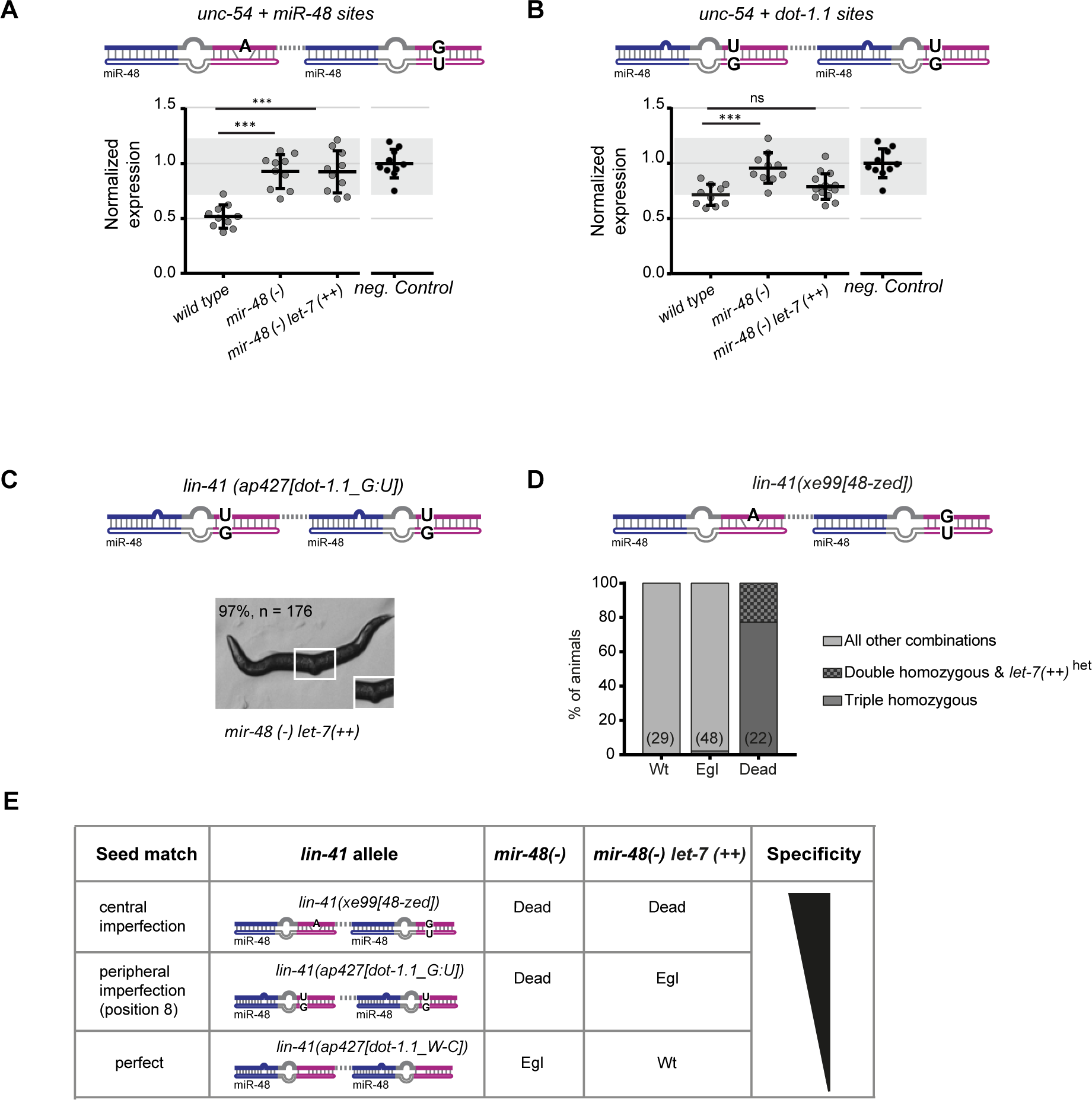
Robust miRNA specificity relies on imperfect seed matches. A), B) Reporter gene expression quantification as in Figure 3B, from which the negative control data are also replotted for reference. A) *unc-54 + miR-48 site* reporter and B) *unc-54 + dot-1.1 site* reporter are assayed in worms of the indicated genotypes. Horizontal line and error bars indicate mean values per condition ± SD,*P < 0.05 and ***P < 0.001, two-tailed unpaired t-test. C) Animals of the *lin-41(ap427[dot-1.1_G:U]), mir-48(-) let-7(++)* genotype are viable but Egl. D) Progeny derived from a cross of *lin-41(xe99[48-zed])* with *mir-48(-) let-7(++)* animals were categorized by phenotype and genotyped, n = 99. *lin-41; mir-48* double mutant animals were in the ‘dead’ phenotype class and even when *let-7* was over-expressed (triple homozygous); only one triple homozygous Egl animal was observed. E) Summary of the effect that different site architectures and miRNA abundance have on silencing *lin-41* alleles “recoded” towards miR-48. *mir-48(-): mir-48(n4097)V; unc-54(CTRL):* wild-type *unc-54 3’UTR; let-7(++): let-7* over-expression allele (MosSCI, V).

To confirm this result on a functional level, we tested whether *let-7* overexpression could suppress the dependence on miR-48 of animals carrying *lin-41* alleles analogous to those in the miR-48-specific reporters, namely the *lin-41*(*ap427[dot-1.1_G:U]*) allele and the newly generated *lin-41(xe99[48-zed])* allele (Figure 4C and 4D, respectively). As predicted by the reporter assay, overexpression of *let-7* rendered *lin-41*(*ap427*); *mir-48(-)* double mutant animals viable, although Egl (Figure 4C). By contrast, we were unable to obtain viable animals of the *lin-41(xe99[48-zed])I; mir-48(-) let-7 (++)V* genotype (Figure 4D). Instead, we readily observed dead animals, which had burst through the vulva. The genotyping revealed that such animals were homozygous for the alleles of interest, *lin-41(48-zed); mir-48(-) let-7(++)* (Figure 4D). [Note that *mir-48(-)* and *let-7(++)* are linked loci on chromosome V, explaining why we did not find dead animals that were *lin-41(48-zed); mir-48(-)* double mutant but lacked the *let-7* over-expression transgene]. In contrast, randomly selected wild-type were never doubly homozygous for *lin-41(48-zed)* and *mir-48(-)*, irrespective of *let-7* transgene status, and only one Egl animal was found *lin-41(48-zed); mir-48(-) let-7(++)* mutant. Hence, although an increase in *let-7* levels can overcome the specificity to miR-48 imposed by seed-distal matches in combination with a perfect seed (Figure 3J) or in the presence of peripheral seed mismatches (Figure 4C), it cannot do so with a central seed bulge or wobble (Figure 4D), at least within the physiological ranges of the expression levels that we tested. We conclude that specificity arises through seed-distal pairing of a miRNA, but that it is enhanced in extent and robustness by appropriate seed match architecture (Figure 4E).

### Loss of miRNA specificity impairs robust development

Our results suggest that sites with central imperfect seed matches, such as those in the *lin-41* 3’UTR, are extremely specific to one miRNA, even when a paralogue is highly expressed. We suspected that such robust specificity would be physiologically relevant in the case of *lin-41*. This is because the *let-7* sisters are all expressed prior to *let-7*, in the L2 stage (Vadla et al., 2012). Given their overlapping spatial expression patterns, lack of mechanisms to prevent *let-7* sisters action on *lin-41* might cause inappropriately early repression of *lin-41*, as speculated earlier (Bartel, 2009; Brennecke et al., 2005). Consistent with this notion, we found that the reporter with the perfect seed match sites “*unc-54 + let-7sites_perfect*” was precociously repressed during the L3 stage, whereas the “*unc-54 + let-7 sites”* reporter was still expressed at the same stage (Figure 5A).

**Figure 5.**
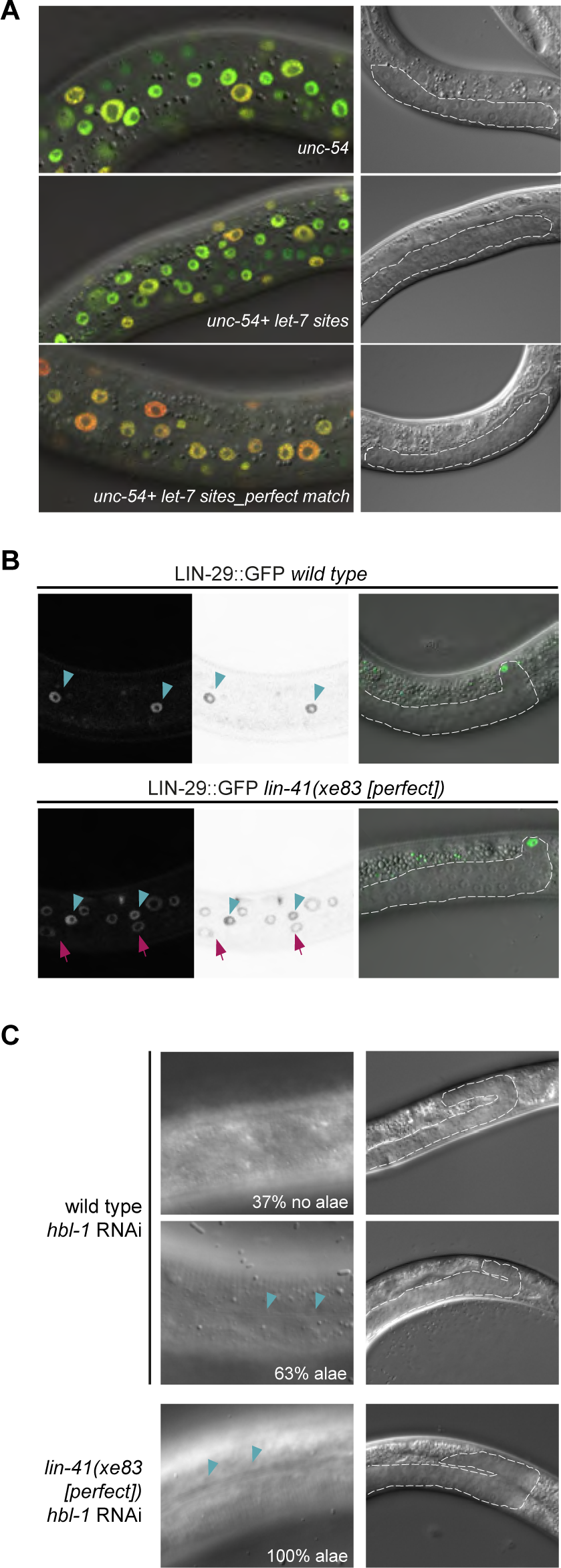
Developmental robustness requires an imperfect *let-7* seed match in *lin-41*. A) Representative confocal images of animals carrying the indicated *unc-54*-derived reporter genes with different *let-7* sites. Animals are at the L3 stage, when *let-7* levels are low but miR-48 levels already high. B) Microscopy images of late L3 worms expressing endogenously tagged LIN-29::GFP (*xe61*) (Aeschimann et al., 2017) in wild-type and *lin-41(xe83[perfect])* background. LIN-29 is detected in hyp7 cells (arrows) only in *lin-41(xe83[perfect])* animals; expression in seam cells (arrowheads) is unchanged. Images in the middle are inverted to increase clarity. C) Representative images of wild type or *lin-41(xe83[perfect])* animals treated with *hbl-1* RNAi. At the L3/L4 transition, 63% (n= 27) of wild type animals have secreted alae, while all the mutant animals do (100%, n = 36). 40x magnification. Gonads of each worm are shown and outlined to confirm appropriate staging.

To test whether this precocious repression of *lin-41* had physiological consequences, we examined the accumulation of LIN-29A, a target of LIN-41. In wild-type animals, LIN-41 translationally represses LIN-29A until the L4 stage, when repression is released following *let-7* accumulation and consequent LIN-41 downregulation (Aeschimann et al., 2017). Premature loss of LIN-41 activity causes inappropriately early activation of LIN-29A and thereby precocious execution of the so-called larval-to-adult transition, which includes fusion of hypodermal seam cells into a syncytium and secretion of an adult cuticular structure termed alae (Slack et al., 2000). We observed LIN-29A levels through use of a *lin-29(xe61[lin-29::gfp::3xflag])* strain in which the endogenous *lin-29* locus has been edited to produce GFP-tagged LIN-29A and B isoforms, and in which loss of *lin-41* activity yields a specific upregulation of only LIN-29A (Aeschimann et al., 2017). At mid-L3 larval stage, wild type animals (staged by the position of the distal tip cell and gonad length) have LIN-29::GFP signal only in their seam cells (Figure 5B). By contrast, animals carrying the *lin-41(xe83[perfect])* allele show additional GFP expression in the major hypodermal syncytium, hyp7, at the same developmental stage (Figure 5B). Therefore, precocious downregulation of *lin-41(xe83[perfect])* is responsible for premature LIN-29 translation and accumulation in the hypodermis, as described for other *lin-41* loss of function alleles (Slack et al., 2000).

Although the *lin-41(xe83[perfect])* animals looked superficially wild-type, the premature upregulation of LIN-29 was sufficient to promote precocious larval-to-adult transition in a sensitized background. Depletion of HBL-1, a transcription factor that is thought to promote larval-to-adult transition in parallel to LIN-41 (Abrahante et al., 2003; Lin et al., 2003) causes partially penetrant and partially expressive precocious heterochronic phenotypes, Figure 5C). By contrast, HBL-1 depletion in *lin-41(xe83[perfect])* animals caused fully penetrant precocious secretion of alae (although weak or patched in some cases) (Figure 5C).

We conclude that loss of specificity of repression by *let-7* alone in the *lin-41(xe83[perfect])* background impairs the robustness of temporal patterning through premature LIN-29 accumulation.

## DISCUSSION

It has been an open question to what extent and by which mechanisms miRNA family members can function non-redundantly despite a shared seed sequence. Previously, it was proposed that redundancy was the rule (Bartel, 2009). Rare occasions of non-redundant function were hypothesized to require targets with both an imperfect seed match and extensive seed-distal pairing to only one specific family member (Brennecke et al., 2005). According to this view, the seed match imperfection impairs silencing by all family members but extensive seed-distal pairing can compensate to facilitate silencing by an individual family member. However, the hypothesis has remained untested, and recent observations have challenged it by providing evidence that non-redundant target binding appears wide-spread and that seed-distal pairing may suffice to achieve specificity (Broughton et al., 2016; Moore et al., 2015).

Our systematic study of *let-7* binding sites on *lin-41* through gene editing and fluorescent reporter analysis with cell-type resolution resolves the discrepant views on specificity-promoting features: We demonstrate that extensive seed-distal pairing to a specific family member suffices to generate a weak but consistent preference for silencing by this family member, but that the extent and robustness of this specificity are low. More robust discrimination among miRNA family requires an imperfect seed match such as conferred by a central bulge or G:U wobble base pair as in the *lin-41* 3’UTR, or a peripheral G:U wobble base pair, as in the *dot-1.1* 3’UTR. Based on this collective evidence, we propose that intra-family specificity is established through different degrees of target complementarity to both the seed and the seed-distal sequences of a miRNA.

Imperfect seed matches can thus be dispensable for specificity, but the effect depends on miRNA levels: A moderate increase in *let-7* levels (~2 fold) could overcome the specificity of a binding site that was specific to miR-48 and had a perfect seed match. However, it only partially did so when the seed match contained a peripheral G:U wobble, and it was insufficient to override sequence-determined specificity when a site contained a centrally imperfect seed match. This implies that *in vivo*, miRNA binding sites are sensitive to miRNA levels and that the seed match quality determines the extent of such sensitivity (Figure 4E).

Given this compelling *in vivo* evidence for an important role of seed mismatches in enhancing miRNA specificity, it is surprising that our computational analysis of the chimera data failed to yield evidence in support of such mismatches as a major criterion for specificity. We consider three possible explanations. First, on a technical level, specificity – as observed in the whole worm by Ago iCLIP chimeric reads – may be driven mostly by differences in miRNA expression patterns rather than intrinsic differences of target binding activity. If only one miRNA family member is co-expressed with a given target, specific chimera will occur, irrespective of seed quality. Because we lack *C. elegans* data on spatial expression patterns of mature miRNAs, we cannot test this possibility. Secondly, the differences between the computational analysis and the experimental dissection of *let-7* family specificity may reflect intrinsic differences in binding promiscuity among miRNA families; i.e., some miRNAs might exhibit promiscuous binding in the presence of a perfect seed and extensive seed-distal matches, and thus require imperfect seed matches to enhance specificity, whereas a majority would not. Consistent with this notion, when we repeated the computational analysis on the levels of individual miRNA families rather than all families in aggregate, we found that for the *let-7* family 75.0 % of targets with perfect seed matches (n = 36) were bound by only a single family member, whereas this fraction was increased to 94.7 % (n = 38) for targets with imperfect seed matches (Supplementary Figure S4A, C). Similar trends were evident for the miR-72 family, although this family shows more sites that are bound by two miRNAs, even in the presence of an imperfect match, possibly because of the close sequence similarity between family members which extends to the seed-distal region (Supplementary Figure S4B, D). However, for the majority of miRNA families we could not recover enough sites from the Ago iCLIP to conclusively show differences. Thirdly, miRNAs might differ in their requirements to achieve specificity not because of intrinsic features but because of their abundance. Although speculative, such a scenario would be compatible with our demonstration that the specificity of *let-7* is an inverse function of its concentration in the absence of major seed match imperfections.

Although the full extent to which miRNA families utilize imperfect seed matches to achieve specific function *in vivo* still remains to be determined, its physiological importance appears evident from the *let-7*:*lin-41* miRNA:target pair, where introduction of a perfect seed match causes loss of specificity and in turn decreased developmental robustness. More generally, our finding that miRNA specificity and functionality rely on miRNA concentrations has major implications for target validation, which has continued to rely extensively on ectopic miRNA expression and a binary yes/no regulation read-out. Our study suggests a much more complex and context-dependent solution as previously hypothesized (Didiano and Hobert, 2006) and consistent with a scenario entertained in the early days of the miRNA field but subsequently disfavored: (Bartel and Chen, 2004) proposed a rheostat model in which the gene silencing activity of a given miRNA is adjusted by two features, namely target site quality, determined by the extent of complementarity to the miRNA, and miRNA abundance. With little explicit experimental testing of such context-dependent function (Doench and Sharp, 2004), and the rising popularity of the “seed-match only” model, the idea has faded from view. However, given the data that we present here, we propose that it is time to revisit this model and subject it to further testing. Certainly, if the goal of target validation is to provide insights into pathway biology, physiology and/or pathology, our results strongly suggest that such validation needs to be conducted in a relevant physiological context and, ideally, involve functional studies such as those offered by direct manipulation of individual miRNA/target interaction through genome editing.

## AUTHOR CONTRIBUTIONS

G.B. conceived the project; designed, performed, and analyzed all the experiments. S.C. performed the analysis of the Ago iCLIP chimeric reads. H.G. conceived the project, designed and analyzed experiments. G.B. and H.G. wrote the manuscript.

## ACKNOWLEDGEMENTS

We thank Kathrin Kunzer and Lan Xu for their help with *C. elegans* strain generation. We are grateful to Matyas Ecsedi for initial observations on target specificity of *let-7* family members. We thank Florian Aeschimann for reagents and helpful discussions, Iskra Katic for worm injections and reagents. We thank Laurent Gelman and Steven Burke for help with confocal imaging; Dr. Roland Nitschke (Life Imaging Center, University of Freiburg, Germany) and Carl Zeiss (Jena, Germany) for sharing the macro for Multiple Position/Tile Imaging acquisitions; Raphael Thierry, Jan Eglinger and Moritz Kirschmann (University of Zurich) for help with image analysis; and Amy Pasquinelli for *C. elegans* strains. Some strains were provided by the Caenorhabditis Genetics Center (CGC), which is funded by the NIH Office of Research Infrastructure Programs (P40 OD010440). We thank Matyas Ecsedi, Iskra Katic and Witold Filipowicz for a critical reading of the manuscript. The work was partly supported by the NCCR RNA & Disease funded by the Swiss National Science Foundation, and the Novartis Research Foundation through the FMI (to H.G.). G.B. was supported by a Boehringer Ingelheim Fonds PhD Fellowship.

## MATERIALS AND METHODS

### Worm handling and strains

Worms were grown using standard methods at 25 °C. The transgenic *unc-54 + miRNA sites* reporter strains were obtained by single-copy integration into the *hTi5605* locus on chromosome II (Frokjaer-Jensen et al., 2012). Injected plasmids were cloned using the MultiSite Gateway Technology (Thermo Fisher Scientific) and the destination vector pCFJ150 (Frokjaer-Jensen et al., 2008) or Gibson assembly (Gibson et al., 2009). All strains are listed in the “Worm Strains” table.

### *unc-54 + miRNA sites* reporters

All *unc-54 + miRNA sites* reporters were constructed using the MultiSite Gateway Technology (Thermo Fisher Scientific) and the destination vector pCFJ150 (Frokjaer-Jensen et al., 2008) or Gibson assembly (Gibson et al., 2009). First, the pGB0 vector was obtained via site-directed mutagenesis (Zheng et al., 2004) of the pDONR P2R-P3_p37 vector to insert the *AscI* restriction site. Then, the pGB01 plasmid was obtained via LR reaction (Gateway LR Clonase II Enzyme mix, Thermo Fisher Scientific; 11791020) of the three entry vectors pdpy-30 x pGFP::H2B x pGB0 and the pCFJ150 backbone.

All the plasmids listed in the Plasmids table were obtained via Gibson assembly of the digested pGB01 plasmid and gBlocks^®^ Gene Fragments (Integrated DNA Technologies) listed below. All plasmids were verified by sequencing. Transgenic worms were obtained by single-copy integration into the *hTi5605* locus on chromosome II, following the published protocol for injection with low DNA concentration (Frokjaer-Jensen et al., 2012). To get a brighter and more physiological mCherry transgene than in (Ecsedi et al., 2015), we exchanged the previous red reporter with a *Pdpy-30::mCherry::H2B::unc-54* transgene integrated on chromosome I and obtained the strain HW1454.

### Genome editing

Mutations in the endogenous *lin-41* 3’UTR sequence were obtained by CRISPR-Cas9 to generate the *lin-41(xe83[perfect]), lin-41(xe76[ap427_W-C])*, and *lin-41(xe99[48-zed])* alleles. Wild-type worms were injected as described in (Katic et al., 2015) with a mix containing 50 ng/μl pIK155, 100 ng/μl of each pGB48 and plin-41sgRNA, 20 ng/ µl repair oligo (see table), *dpy-10* co-crispr mix containing 100 ng/ml pIK208 (Addgene plasmid #65630) and 20 ng/ml AF-ZF-827 oligo PAGE purified (IDT). Single F1 roller progeny of injected wild-type worms were picked to individual plates and the F2 progeny screened for the mutated allele using PCR assays and sequencing. The alleles were outcrossed three times to the wild-type strain.

#### *let-7* over-expression

A *let-7(++)* strain (HW 1909 [*xeSi287*, V]) was obtained by injection of the plasmid pGB26, obtained via Gibson assembly (Gibson et al., 2009) of the PCR amplified minimal rescue fragment from (Reinhart et al., 2000) and the pIK37 plasmid. Transgenic worms were obtained by single-copy integration into the *oxTi365* locus on chromosome V (universal MosSCI strain #EG8082 (Frokjaer-Jensen et al., 2014).

### Reporter Quantification

For confocal assays, worms were grown at 25 °C. *Let-7ts* worms were maintained at 15 °C and adults were transferred to 25 °C 48h before imaging. Z-stacks of 0.313 µm µm thickness were acquired in green, red and transmitted light channels at 40x magnification on a Zeiss LSM700 confocal microscope coupled to Zeiss Zen 2010 sotware equipped with a mul.-position tile scan macro. The z-stacks were stitched together and compiled into a single image using scripts in Matlab and Fiji (Schindelin et al., 2012).

For data analysis, late L4 worms were selected based on visual inspection of gonad length and vulva morphology (Mok et al., 2015). 10-14 vulva cells were selected in the ‘cell counter’ macro in Fiji. Images around these seed points were de-noised using a Richardson-Lucy algorithm and segmented using an Otsu global threshold. Remaining holes were filled using a morphological filter. Signal intensity in the green channel was divided by the red signal intensity for each cell; relative signal intensities were then averaged for each worm. 10-12 vulva cells in 5-10 worms per genotype were quantified, mean signal intensity and SD were calculated and graphed using GraphPad *Prism sojware*.

### Confocal analysis of LIN-29 precocious accumulation

Synchronized arrested L1 larvae of animals carrying endogenously tagged LIN-29, *lin-29(xe61[lin-29::gfp::3xflag])*, in wild type or *lin-41(xe83[perfect])* background, were plated on food and incubated at 25 °C on 2% NGM agar plates with *Escherichia coli* OP50 bacteria and imaged at the L3 stage (20-22h ater plating). Images were acquired in green and transmitted light channels (with Differential Interference Contrast, DIC) with 40x/1.3 oil immersion objective on a Zeiss LSM700 confocal microscope coupled to Zeiss Zen 2010 sotware. Further image processing was performed with Fiji (Schindelin et al., 2012).

### Ago iCLIP reads analysis

Reproducible miRNA-target sites were extracted from supplementary table S2 from (Broughton et al., 2016). Target sequences were retrieved from the UCSC October 2010 (ce10) genome assembly (Rosenbloom et al., 2015), using the BSgenome.Celegans.UCSC.ce10 package in R. Sites in 3’ UTRs or CDSs were iden.fied by intersec.ng all sites with annotated 3’ UTRs or CDSs from the ce10 Ensembl gene annota.ons. MicroRNA family informa.on, including mature miRNA and seed sequences, was downloaded from TargetScan release 6.2 (Agarwal et al., 2015). Chimeras were predicted by calling RNAhybrid on all miRNA-target pairs using the command ‘RNAhybrid -b 1 -c -s 3utr_worm’ (Rehmsmeier et al., 2004). Perfect sites were identified by searching for an exact match to the corresponding seed in the predicted bound miRNA and target sequences for each chimera. Imperfect sites were defined as any site containing a single bulged nucleotide, a single G:U wobble or a single mismatch in the target between positions 2-8.

The number of paired 3’ nucleotides for each chimera was determined by counting the number of nucleotides in the mature miRNA predicted to be bound by RNAhybrid downstream of the seed match. For imperfect sites where an exact seed match was not present, 3’ paired nucleotides were considered as any predicted to be bound downstream of the 7^th^ paired nucleotide ater trimming an initial U if present (corresponding to position 8 of the seed).

RNA-seq data from *alg-1(-)* vs. WT were downloaded from the SRA (accession number: SRP078368). Reads were aligned against the ce10 genome assembly using the qAlign function from the QuasR R package (Gaidatzis et al., 2015) with default se„ngs. Reads were counted in all annotated transcripts from the ce10 Ensembl gene annotations. Counts were normalized by the mean number of reads in transcripts in both libraries, and a log2 fold-change for each transcript was calculated between *alg-1(-)* and WT. Transcripts were considered to be targets of a particular miRNA if a corresponding miRNA-target site was found in either the 3’ UTR or CDS, respectively. The empirical cumulative distribution of the log fold-change for each class of sites was calculated using the ecdf function in R.

Minimum free energy predictions were taken directly from RNAhybrid. All computations were performed using R (version 3.4.0) in the RStudio environment (version 1.0.143).

### Worm strains

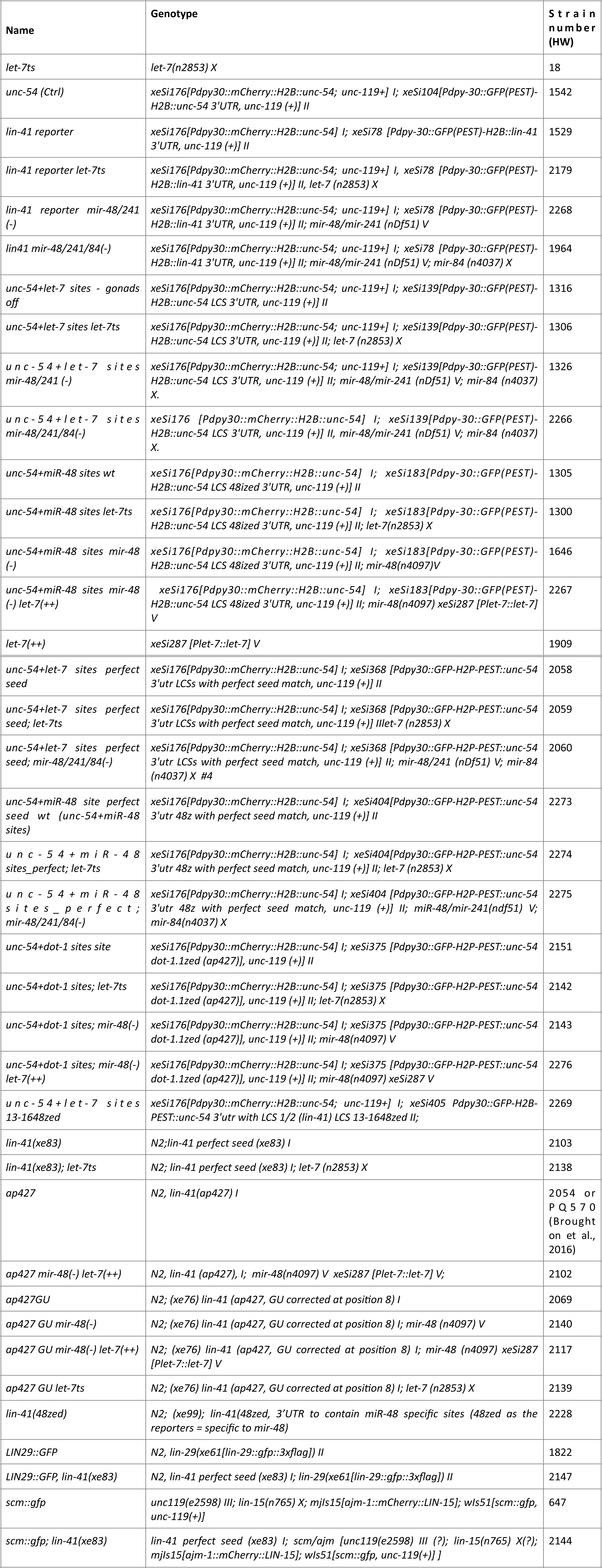

### Plasmids

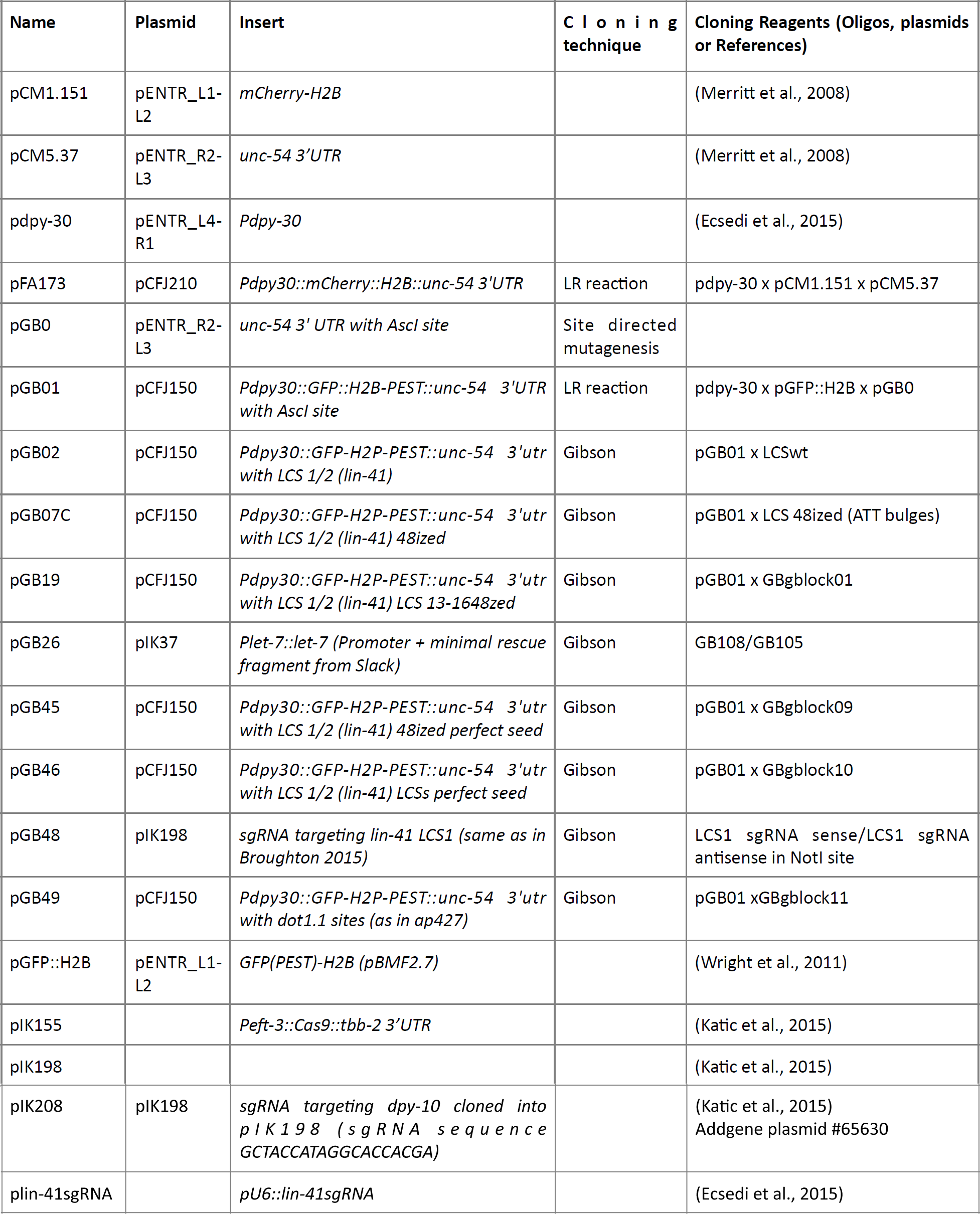

### Primers

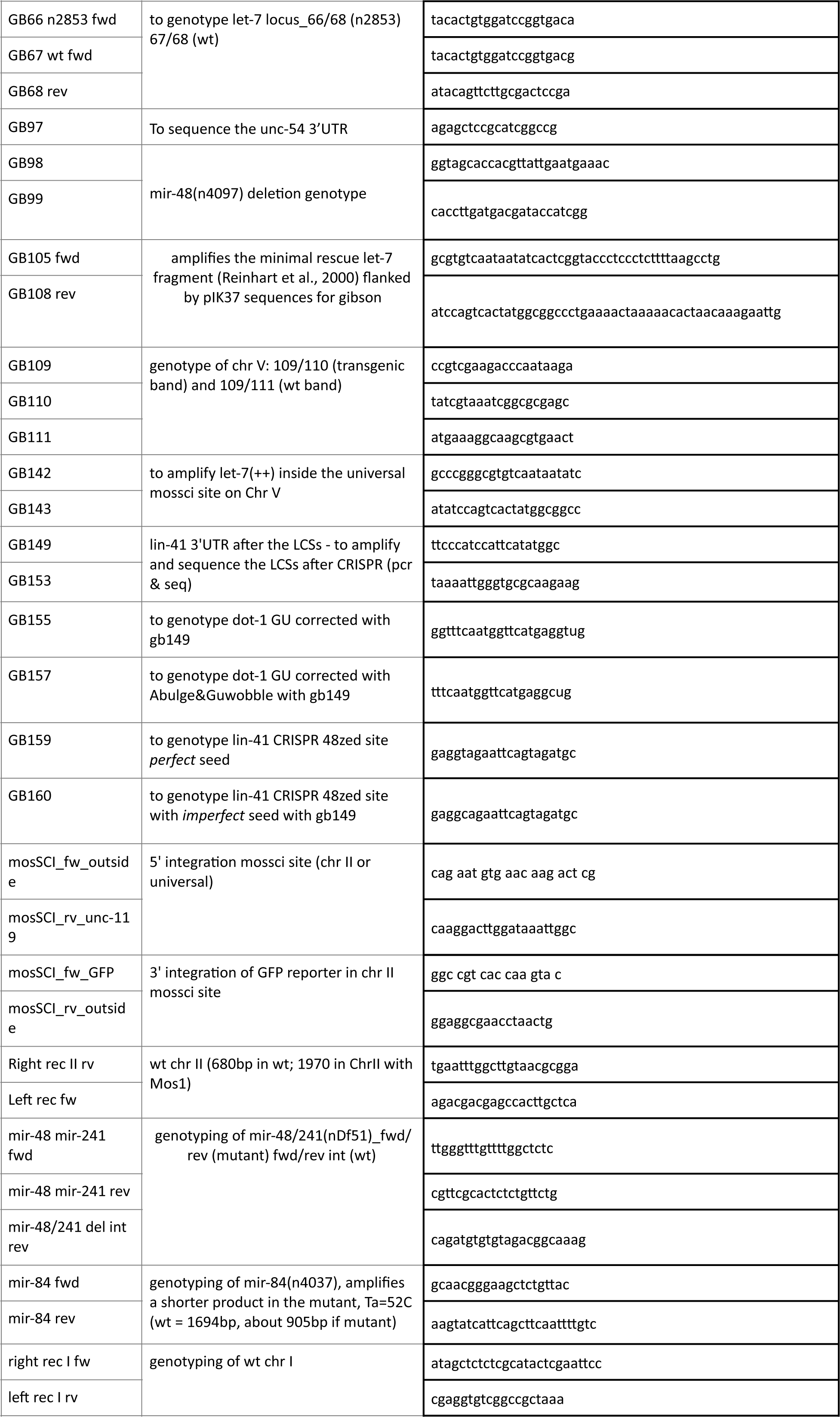

### gBlocks and homologous recombination oligos

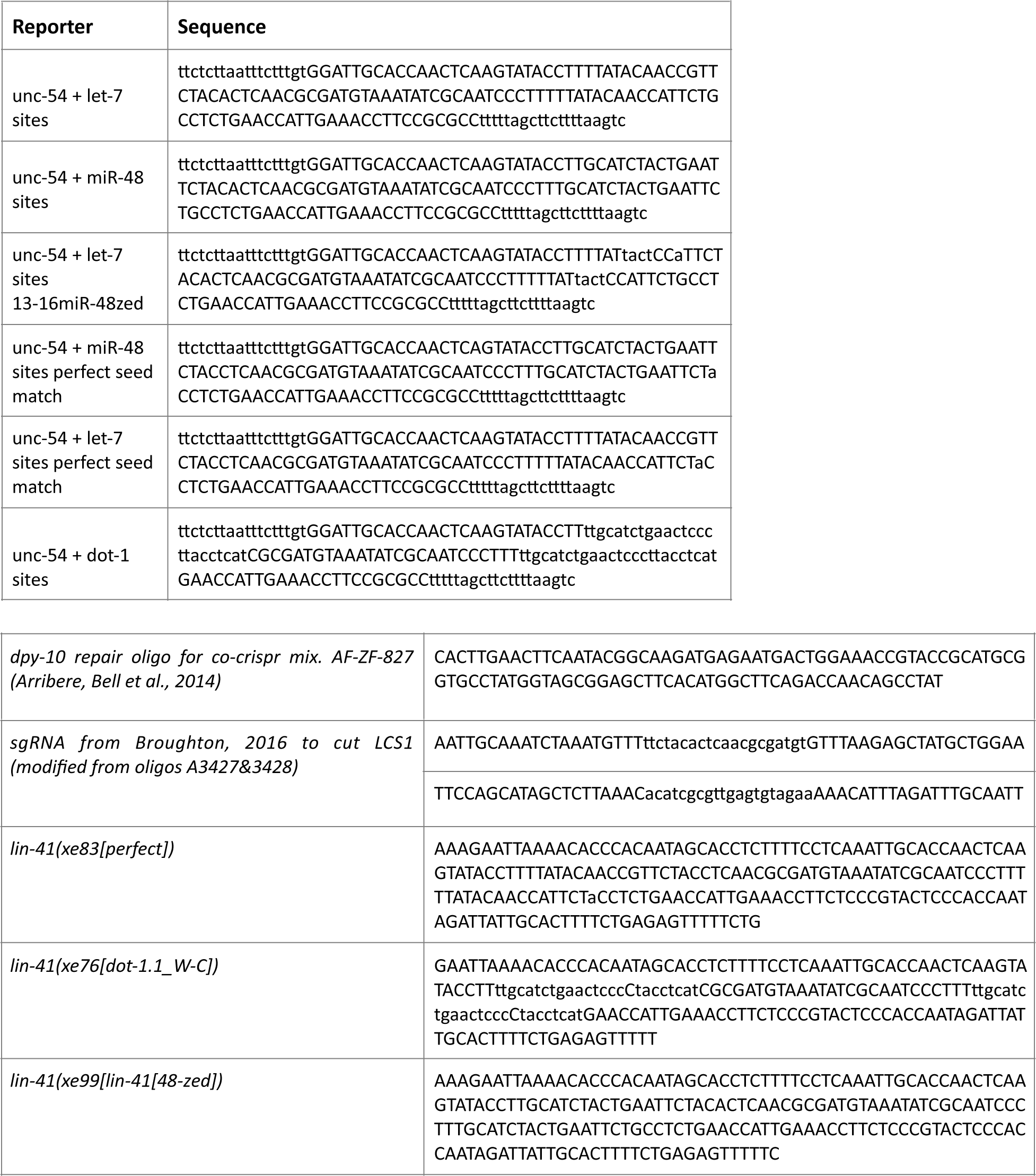

## Supplementary Figures

**Supplementary Figure S1.**
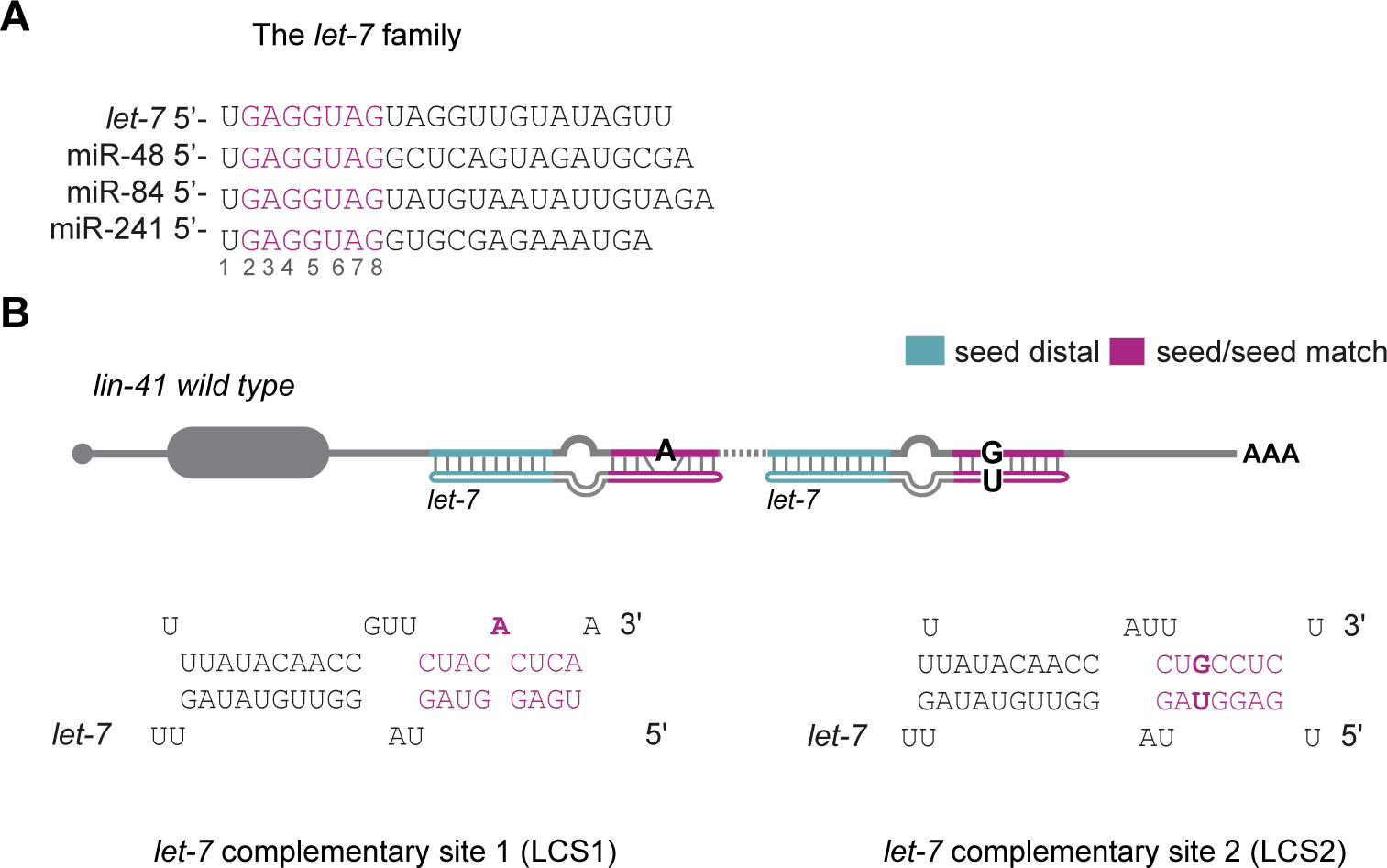
A) The ***let-7*** family sequence with the seed sequence highlighted in magenta B) The two ***let-7*** complementary sites (LCS 1 and LCS2) in the ***lin-41*** 3’UTR of ***C. elegans*.** Each site contains an imperfect seed match (a bulged A and a G: U wobble, in bold) to the ***let-7*** family and an extensive seed-distal pairing to ***let-7*** only.

**Supplementary Figure S2.**
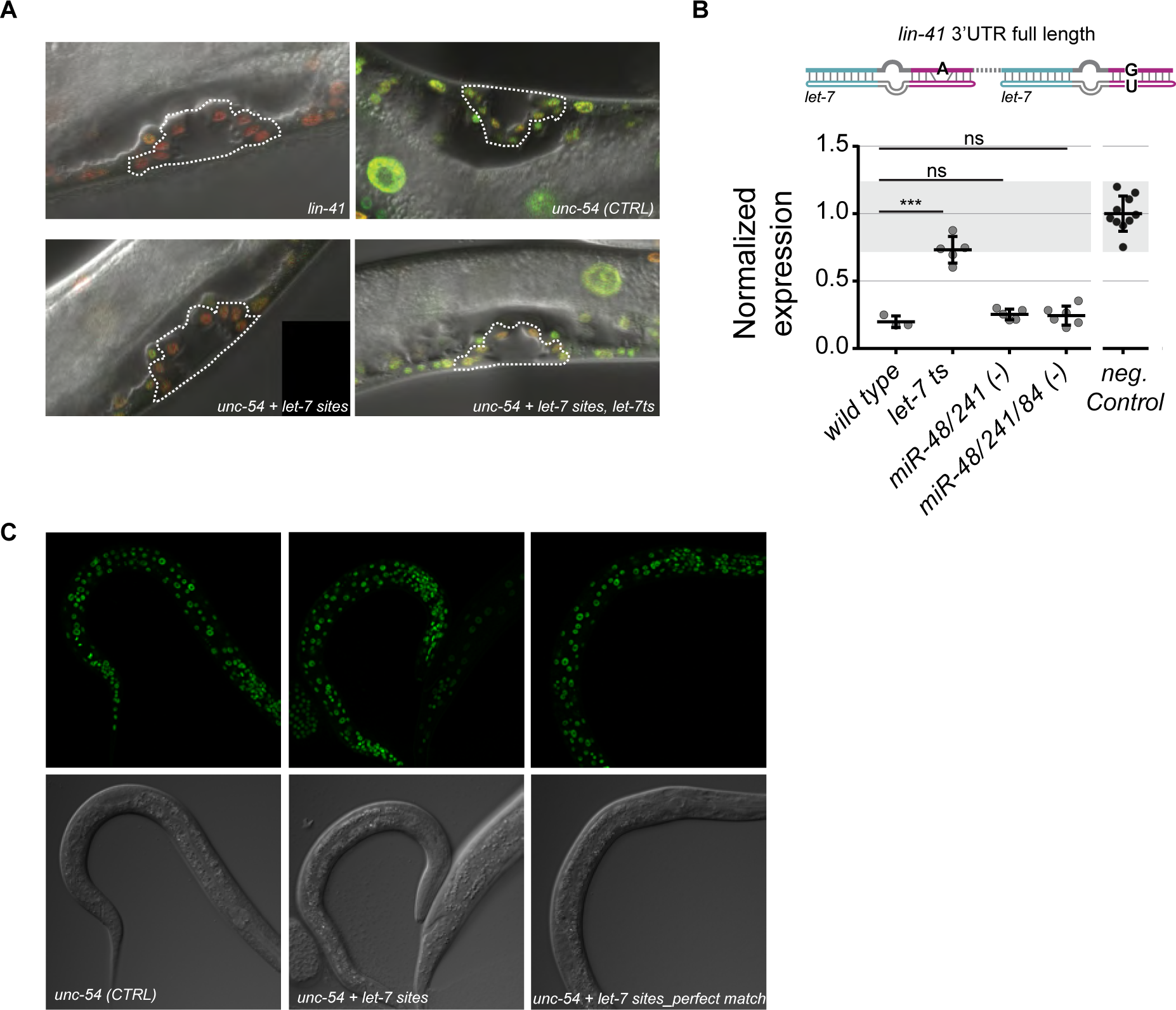
A) Representative confocal images of animals carrying the red mCherry reporter (used for normalization) and GFP reporters with the following 3’UTRs: *lin-41* full length, *unc-54* (Control) and *unc-54 + let-7* sites in wild type and *let-7ts* background. *Unc-54 + let-7* sites is silenced like the *lin-41* reporter, and its silencing depends on *let-7*. Dashed lines outline the vulvae of the animals, which confirm appropriate late Larval stage 4 (Late L4). B) Quantification of the *lin-41* full-length reporter in the vulva cells of late L4 animals. Each dot in the graph represents the average of the GFP signal intensity divided by the mCherry intensity of a single animal per condition. 10-12 vulva cells were quantified per worm. Values are normalized to the average value of the GFP/ mCherry ratio of the *unc-54* reporter, replotted for reference from Figure 3B. Horizontal line and error bars indicate mean values per condition ± SD. *P < 0.05 and ***P < 0.001, two-tailed unpaired t-test. C) Representative confocal images of young L1/L2 worms carrying reporters with the following 3’UTRs: *unc-54*, *unc-54 + let-7* sites and *unc-54 + let-7sites_perfect match* in wild type animals. None of the reporters are silenced in young larvae. *Unc-54* reporter = control. Top panels: GFP & mCherry overlay; bottom panels: DIC. *Let-7ts: let-7(n2853) X*, temperature sensitive lesion, grown at the restrictive temperature 25C; *mir-48/mir-241/ mir-84 (-): (nDf51) V, (n4037) X; mir-48(-): mir-48(n4097)V; unc-54(CTRL):* wild-type *unc-54 3’UTR*.

**Supplementary Figure S3.**
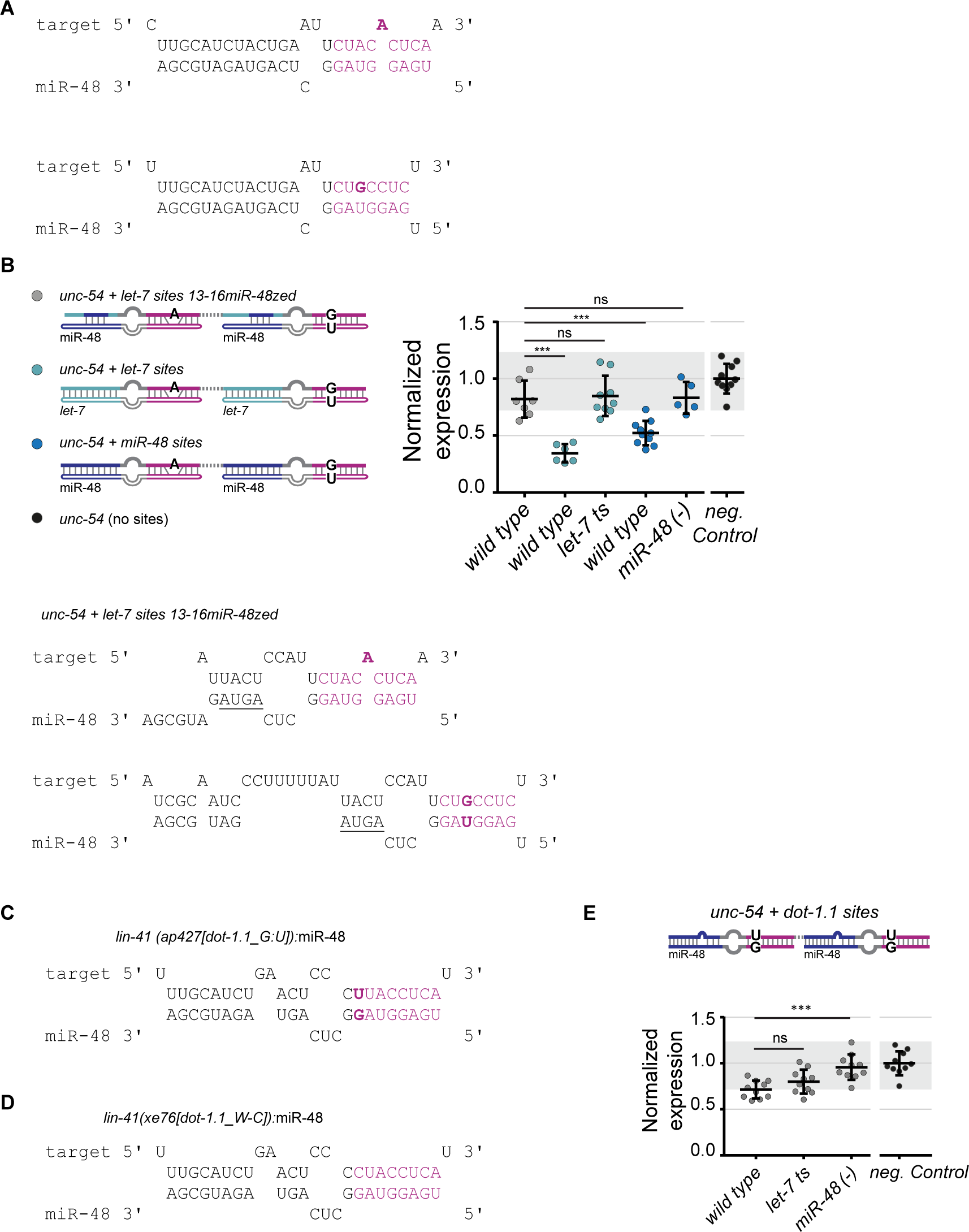
A) Schematic of the duplexes of the binding sites in the *unc-54 +* miR-48 reporter paired to miR-48 as predicted by RNAhybrid (Kruger and Rehmsmeier, 2006). B) Quantification of a reporter containing *let-7* sites modified to pair miR-48 at position 13-16. The reporter fails to be silenced in wild type animals. The reporters Unc-54 + *let-7* sites and *unc-54* + miR-48 sites in wild type and *let-7ts* or *mir-48(-)* background, respectively, are reported as reference from Figure 3B and 3D, neg. control from Figure 3B. Horizontal line and error bars indicate mean values per condition and ± SD. *P < 0.05 and ***P < 0.001, two-tailed unpaired t-test. C),D) Schematic of the duplexes of the binding sites in the *lin-41(ap427[dot-1.1_G:U]) and lin-41(xe76[dot-1.1_W:C])* paired to miR-48 as predicted by RNAhybrid (Kruger and Rehmsmeier, 2006). E) Quantification of the reporter “*unc-54 + dot-1.1 sites*” reveals miR-48 specificity in the vulva*. let-7ts*: (*n2853) X*, temperature sensitive lesion, grown at the restrictive temperature 25C; *mir-48(-)*: *mir-48(n4097*)V; Neg. Control reported from Figure 3C. Mean ± SD. *P < 0.05 and ***P < 0.001, two-tailed unpaired t-test.

**Supplementary Figure S4.**
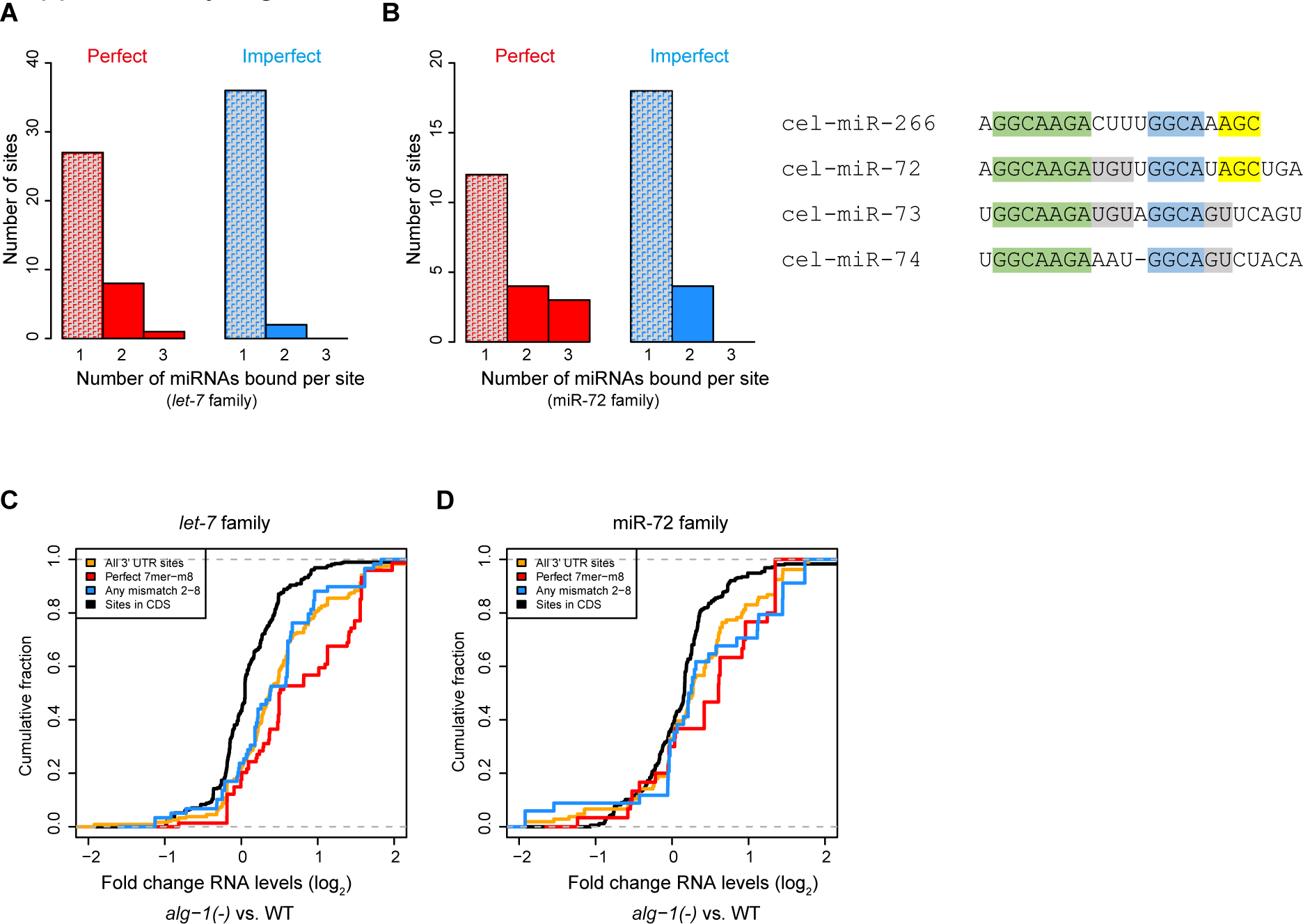
A) Abundance of miRNA sites located in 3’UTRs found in ALG-1-chimeras bound by a miRNA belonging to the *let-7* family and containing either a perfect seed match or an imperfect seed match (one mismatch in the seed/seed match duplex (nucleotides 2-8: bulges, wobbles or any mismatch) B) Same as in (A) but for chimeras containing a miR-72 family member C), D) Upregulation in *alg-1(-)* of transcripts containing sites for a member of the *let-7* (c) or miR-72 (d) family. Sites with imperfect seed match (blue) or a canonical 7mer-m8 (red) compared to sites in CDS (black) or any miRNA binding site located in 3’UTRs.

